# Comparative analysis of machine learning algorithms on the microbial strain-specific AMP prediction

**DOI:** 10.1101/2022.01.28.478081

**Authors:** Boris Vishnepolsky, Maya Grigolava, Grigol Managadze, Andrei Gabrielian, Alex Rosenthal, Darrell E. Hurt, Michael Tartakovsky, Malak Pirtskhalava

**Author notes:** Corresponding Authors: B. Vishnepolsky., M. Pirtskhalava.

## Abstract

The evolution of drug-resistant pathogenic microbial species is a major global health concern. Naturally occurring, antimicrobial peptides (AMPs) are considered promising candidates to address antibiotic resistance problems. A variety of computational methods have been developed to accurately predict AMPs. The majority of such methods are not microbial strain-specific (MSS): they can predict whether a given peptide is active against some microbe, but cannot accurately calculate whether such peptide would be active against a particular microbial strain. Due to insufficient data on most microbial strains, only a few MSS predictive models have been developed so far. To overcome this problem, we developed a novel approach that allows to improve MSS predictive models (MSSPM), based on properties, computed for AMP sequences and characteristics of genomes, computed for target microbial strains. New models can perform predictions of AMPs for microbial strains that do not have data on peptides tested on them. We tested various types of feature engineering as well as different machine learning (ML) algorithms to compare the predictive abilities of resulting models. Among the ML algorithms, Random Forest and AdaBoost performed best. By using genome characteristics as additional features, the performance for all models increased significantly—on average by 7%—relative to models relying on AMP sequence-based properties only. Our novel MSS AMP predictor is freely accessible as part of DBAASP database resource at https://dbaasp.org/tools?page=genome-prediction

## 1. INTRODUCTION

One class of molecules that are considered as promising candidates to treat the problems linked with antibiotic resistance are antimicrobial peptides (AMPs). Over the years, many computational methods, including machine learning and artificial intelligence, were trained on diverse AMP datasets, and evaluated for their performance. [1]. The importance of large, well-balanced datasets, as well as powerful computational methods and informative descriptors has been reviewed in [1]. Several drawbacks of AMP datasets used in the existing AMP prediction tools are discussed in [2]. Understanding what physico-chemical and statistical descriptors (features) of peptides are essential for including in the analysis, remains critical both for better understanding of AMP’s mechanisms of action and improving the resulting models prediction quality. AMP prediction algorithms used a variety of machine learning methods, including support vector machines (SVM), fuzzy K-nearest neighbours (FKNN), random forest (RF), neural networks (NN). Based on these methods, various online prediction services have become available. Some of these tools allow for simultaneous analysis using various machine learning approaches for AMP prediction, which becomes especially significant when all models agree on a prediction [3,4]. To improve the performance for non-consensus cases, the ensemble models have been proposed [5,6], whose predictions are based on combining the outputs of different models (such as SVM and random forest), weighed by derived parameters. However, given the scarcity of data on AMPs, such composite models may have a tendency to overfit the training set, with the questionable performance outside of it.

Despite the improvements in predictive algorithms, most methods share the same conceptual issues. First, during the model development, they do not take into account the information on target strains, although the antimicrobial potency of AMPs strongly varies on strain-specific bacterial envelope types. Second, for models training, most of the methods use the peptides with incomplete or non-existent experiment-based information on their antimicrobial activities. These drawbacks lead to overall uncertainty in the interpretation of prediction results. For instance, Lee at all. [7] have developed a SVM classifier-based predictive model of AMP, that has shown excellent performance against the blind test set, with a prediction accuracy of 91.9%, specificity of 93.0%, and sensitivity of 90.7%. At the same time, a detailed analysis of the algorithm’s predictions has shown that the features, learned by the support vector machine, do not correlate with antimicrobial activity, but with a peptide’s ability to generate the negative Gaussian membrane curvature. Consequently, the authors have coined their SVM classifier as a general detector of membrane activity in peptide sequences but not as a true predictor of antimicrobial activity. It is worth noting, that training sets used in the development of the majority of available predictive methods use AMPs of arbitrary length. Yan, Bhadra et al. [8] have explored four state-of-the-art AMP prediction methods to test whether they can be used for predicting short-length AMPs. Prediction accuracy, achieved with the help of these models was between 65% and 73%, which is significantly worse than the previously reported accuracy of 90% −95% for models trained on peptides of any length.

The majority of AMP databases do not provide information about antimicrobial activity against particular strains (at least in a convenient-for-analysis form). DBAASP is one exception from this ‘rule’, providing reports about antimicrobial activity against particular strains, extracted from scientific publications. Recognizing this advantage, many strain-specific predictive models have been trained on datasets from DBAASP [9–15]. For example, a method developed by Speck-Planche and coworkers [9,10] uses a multi-target approach and discriminant analysis to predict antimicrobial activity against different bacterial strains. Another MSS AMP prediction method described in [11] uses a random forest algorithm for recognizing antimicrobial activity as well as identifying molecular descriptors underpinning antimicrobial activity of investigated peptides. For some strains, very small amount of data is available and thus the prediction results for these strains may not be reliable. Wang et al. [16], using a bidirectional LSTM recurrent neural network classifier, attempted to design novel AMP sequences with potential activity against *E.coli*. Another example of open access predictions resource is described in [12], where Gull and Mina offer a web server for predicting AMP with activity against particular strains [12]. The authors tried to overcome the problem of insufficient training data by using the target microbe’s genome information along with the sequence-based parameters. This is, to our knowledge, the only algorithm where the authors include the information about target microbes in the model to generate MSS AMP predictions. Compositions of mono, di-, tri- and tetranucleotides have been used as genomic features. Only the amino acid compositions have been used as peptide descriptors for the model. Yet another example is described in [15], where the authors developed model using a Siamese network architecture to learn from pairs of AMPs to predict how their activity differs against 10 different genera of bacteria.

In our previous paper [17], a predictive model for linear AMPs active against *Escherichia coli ATCC 25922* has been described, which relies on a semi-supervised machine-learning approach with a density-based clustering algorithm. Subsequently, prediction models for some other microbial strains were developed as described in [18]. To support and further the involvement of research community in computer-aided studies of AMPs, we implemented our current model as open access, on-line prediction tool, available on DBAASP website.

Using informative descriptors of microbial organisms as features could allow not only to improve the performance of MSS AMP predictive models in general, but to perform predictions for cases when experimental data is insufficient or absent. The main question then becomes, *how should a microbe strain be described to include this feature(s) in the model?* The most convenient description would be using behavior of microbial strain against particular set of AMPs (one example would be using susceptibilities of the microbe for a set of AMPs [17]). However, obtaining a statistically reliable set of peptides that have been tested on a wide spectrum of microbes is currently challenging, to say the least. Therefore, we considered at purely genome-based statistical descriptors that are easily computed for any available genomic sequence.

Genome-based characteristics of microbes can be divided into absolute characteristics (compositions of oligonucleotides in the genomic sequence) and comparative characteristics (describing similarity between genomes). Furthermore, similarity-based characteristics can be divided into 2 classes: the first class includes characteristics that are based on similarity between particular genes which are widely used as phylogenetic markers, such as 16S RNA [19], DNA gyrase [20] and others. The second class includes characteristics which reflect similarity between full genomes. The second class of characteristics reflect (computer-assessed) pairwise genomesequence-based similarity and were coined as the overall genome related index (OGRI) [20]. Average nucleotide identity (ANI) [22] and digital DDH (dDDH) [23] are most widely used OGRI for delineation of bacterial species. To our best knowledge, similarity-based characteristics of genomes have not been previously used as features in training MSS AMP predictive models.

In this work, we perform a comparative analysis to find the ‘best’ combination among the various sets of features and machine learning approaches to generate MSS AMP predictive models. Our comparative analysis has been conducted on special benchmarks, created with the help of the DBAASP data.

## 2. METHODS

### 2.1 Machine learning approaches and algorithms

Six ML algorithms have been chosen from the Weka workbench (https://www.cs.waikato.ac.nz/~ml/weka/) and trained on the DBAASP data in order to reveal the most stable and accurate AMP prediction algorithm. These are: Random Forest, LibSVM, K-nearest neighbours, RealAdaBoost, Multilayer Perceptron (Neuron network), Dl4jMlpClassifier (Deep neural network-based classifier). The models’ default parameters have been used for training, except for RealAdaBoost where as base classifier Random Forest has been chosen. All models have been evaluated with the help of 10-fold cross validation.

### 2.2 Microbial strains

Microbial strains (MS), that have been used to gather information on susceptibilities for AMPs have been chosen based on availability of data in DBAASP [24] and Genbank [25]. MSs for which there is data on genome sequences have been divided into two groups: ones for which data on susceptibility available in DBASSP is enough to build a good predictive model, and ones that don’t have enough data to train a separate predictive model, but present high threat for healthcare [26] — the data from this second group has been combined with the data from the first one to train a genome-based predictive model. Examples of strains that have enough data on interaction with the AMPs are: *Escherichia coli ATCC 25922, Staphylococcus aureus ATCC 25923* and *Pseudomonas aeruginosa ATCC 27853* (Table S1). Strains that are considered as major problem for healthcare and that are included in our training set, are: *Klebsiella pneumoniae ATCC 700603, Salmonella typhimurium ATCC 14028, Enterococcus faecalis ATCC 29212, Acinetobacter baumannii ATCC 19606* and *Bacillus subtilis ATCC 6633*. Majority of the latter group are part of ESKAPE pathogens [27].

### 2.3 Genome-based characteristics of microbial strains

Genome-based characteristics of MS can be divided into absolute and comparative. Mono-, di-, tri- and tetra-nucleotide compositions of the genome are an absolute characteristic, which is often used to assess level of similarity between species [28–33]. They encode the genome by means of 4, 16, 64 and 256-dimensional vectors, respectively. We have used a combination of mono+di, mono+tree and mono+tetra nucleotide compositions as well, because GC content difference has large influence on microbial classification using oligonucleotide frequency [33] and mono nucleotide composition reflects GC content. Such combinations encode genomes using 20, 68 and 260-dimensional vectors, respectively. It is widely believed, that the most effective way to delineate microbial species based on their genomes’ similarity is by making use of comparative characteristics, such as ANI [22] and dDDH [23]. ANI and dDDH are assessed between the pairs of genomes. According to [34], dDDH yielded higher correlations with experimental values of DDH than ANI, and therefore we have used dDDH in our comparative analysis. To encode the strains chosen for assembling the training set (see Table 1), five reference genomes have been selected from GenBank [25]. They are the genomes of *Escherichia coli str. K-12, Staphylococcus aureus subsp. aureus NCTC 8325, Pseudomonas aeruginosa PAO1, Bacillus subtilis subsp. spizizenii ATCC 6633* and *Mycobacterium tuberculosis H37Rv*. The first four species (two Gram- and two Gram+) represent the most-often-used species in experiments for defining antimicrobial activity of peptides and there are large volumes of data for them in the database. Mycobacterium tuberculosis are Gram-indeterminate bacteria with well-studied genomes [35]. Selected strains are presented in GenBank as reference genomes (RG) for the corresponding species and therefore they have been chosen as reference for assessing comparative genome-based attributes. Each strain from table 1 has been encoded using 5 comparative attributes, becoming a 5-dimensional vector whose components dDDH j (j =1,…,5) have been assessed by the Genome-to-Genome Distance Calculator (GGDC) [23].

**Table 1.**
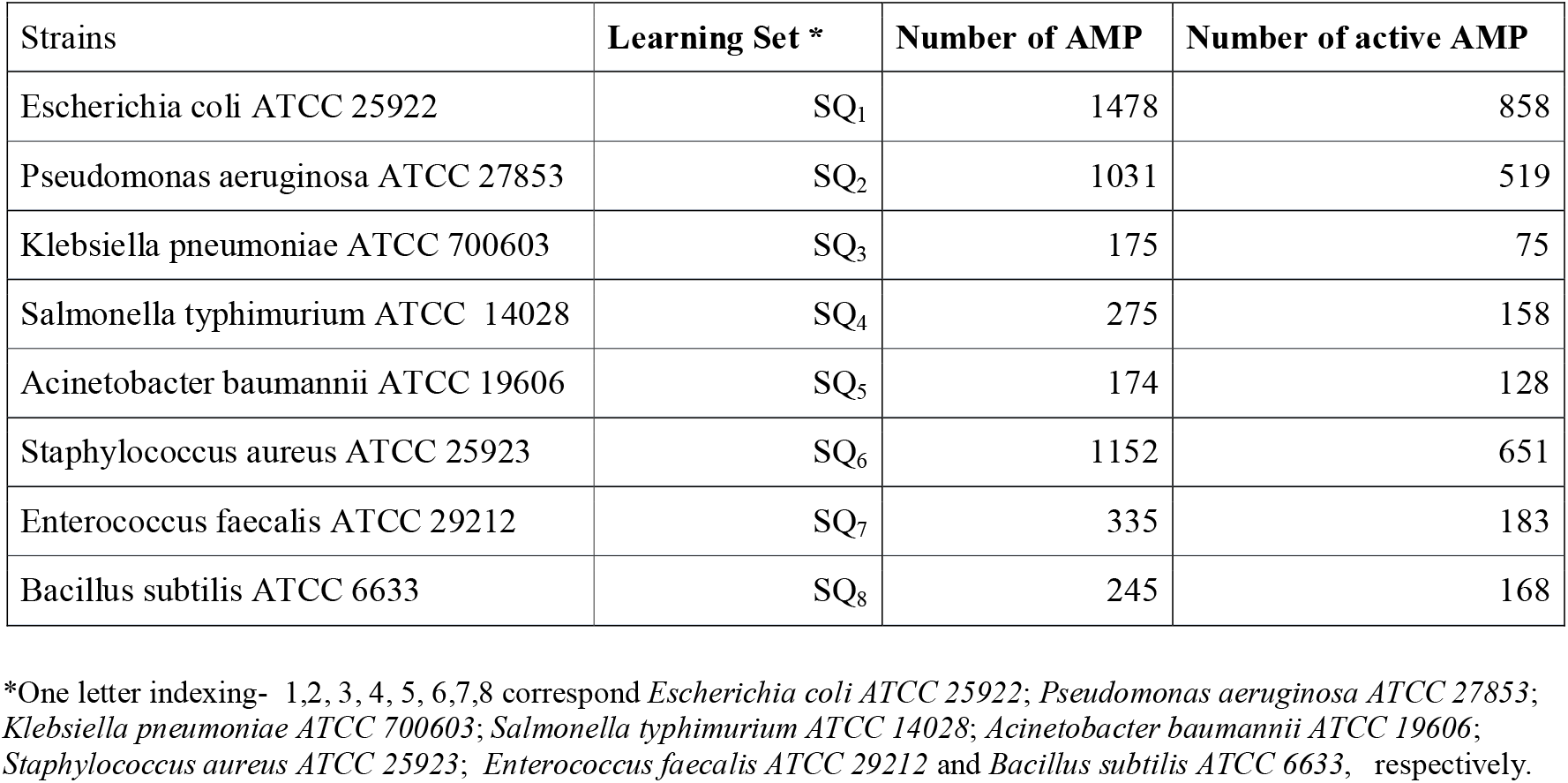
Strains selected for the comparative analysis.

### 2.4 AMPs, chosen for training sets

The following filters have been used to DBAASP database to select peptides for creating training sets: length 9-30, no amino acid modifications, no inter-chain bonds, no N-terminal modifications. The number of peptides included in the training set for each microbial strain is given in Table 1.

To define a peptide’s class, its MIC value has been used. MIC is a widespread measure of the potency of AMP. The threshold for definition of an AMP as “active” is not strictly defined across different resources. For instance, APD database [36] that stores information on active AMPs uses the following criterion: MIC < 100. We used this threshold as a starting point for our own classifications, defining an AMP as *not* active in cases when MIC > 100. To define an AMP as active, we had to take into consideration the peculiarities of the broth microdilution method, which is a major method for the assessment of MIC. Broth dilution method is characterized by large error rates [37]; uncertainty of the assessment can be equal to two dilutions. Therefore, as the criterion of activity we have used the following threshold: MIC < 25. Consequently, our training sets have been created by adopting the following thresholds of antimicrobial activity: MIC < 25 μg/mL for the positive samples and MIC > 100 μg/mL for the negative ones [17].

### 2.5 Sequence-based characteristics of AMPs

The majority of computational approaches for studying AMPs use five major types of features for description of their structures: (1) composition features (2) position features (3) structure features (4) physicochemical properties and (5) similarity features. Moreover, some features may belong to several types. We used the following sequence-based characteristics of AMP: Normalized Hydrophobic Moment (NHM), Normalized Hydrophobicity(NH), Net Charge(NC), Isoelectric Point (IP), Penetration Depth, (PD), Tilt Angle(TA), Linear Moment (LM), Cyclic Linear Moment (CLM), Propensity to in vitro Aggregation (PIA), Propensity to ppII coil (PII), Angle Subtended by the Hydrophobic Residues (AH), Amphiphilicity Index(AI), Propensity to Disordering (PD),: Composition of Arg (CR), Composition of Trp (CW), Composition of Lys (CK), Composition of Pro (CP), Composition of Hys (CH), Composition of Gly (CG), Peptide Length (PL).

The majority of these features (such as NHM, NH, NC, IP, PIA, PL) have been widely used in model development, including our own previous works. A detailed discussion of these features can be found in [38]. PD, TA, AH, AI and PII are features that we have used in AMP predictive model development in the past [24,38]. Using compositions of selected amino acids (such as CR, CW, CK, CP, CG, CH) as features is linked with the statistically reliable abundance of these amino acids in the sequences of AMP, and consequently with their key role in antimicrobial activity [39]. Amphipaticity of AMP together with helical distribution of amino acids can be reached by simple linear segregation of hydrophobic and hydrophilic amino acids along the polypeptide chain. We have therefore used two features to describe linear separation of hydrophobic and hydrophilic residues along the chain. These are Linear Moment (LM) and Cyclic Linear Moment (CLM). Their definition is presented below:

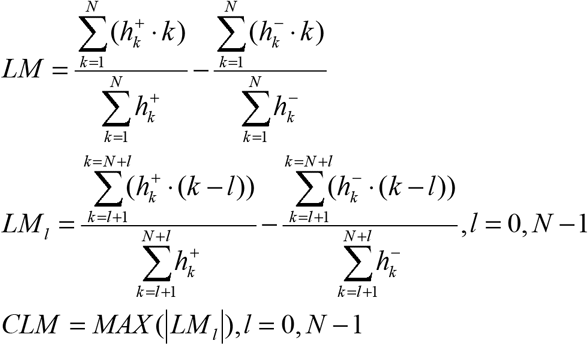

Where *N* - peptide length, 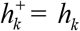 or 0 if *h_k_* ≤0; 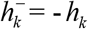 if *h_k_* <0, or if *h_k_* ≥0; *h_k_* - element of the Kyte-Doolittle hydrophobic scale for the *k-th* residue [40]; 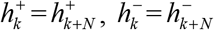

### 2.6 Training sets

#### 2.6.1 Sequence-based training set

For each MS, the sequences of AMPs tested against it have been gathered from DBAASP and a training set of SQ type has been created. 8 trainig sets, SQi (i=1-8), have considered, where i = 1 corresponds to *Escherichia coli ATCC 25922*, i = 2 - *Pseudomonas aeruginosa ATCC 27853*, i = 3 - to *Klebsiella pneumoniae ATCC 700603*, i = 4 - to *Salmonella typhimurium ATCC 14028*, i = 5 - to *Acinetobacter baumannii ATCC 19606*, i = 6 - to *Staphylococcus aureus ATCC 25923,* i = 7 - to *Enterococcus faecalis ATCC 29212*, and i = 8 - to *Bacillus subtilis ATCC 6633*. Such training sets unite instances (vectors of the sequence-based attributes) characterizing two classes of AMPs: active against particular microbial strain (positive set) and non-active against it (negative set).

#### 2.6.2 TS+APS - based training set

To overcome the problem of insufficient data encoding of a pairs of “Target strain - AMP sequence” (TS+APS) has been performed and instances as vectors of attributes characterizing target strain and AMP have been used. To develop TS+APS-based MSSPMs the various genome features (GF) have been used. Correspondingly, the following training sets have been created: SQTS_i j_ (i=1,..8) where index i corresponds to MSs and j corresponds to GFs. Here, i=1,…,8 defines 8 MS in the same correspondence as in the previous paragraph. Index j=1,..,9 defines GFs. The values of this index from 1 to 7 corresponds to oligonucleotide compositions of genome: mono, di, tri, tetra, mono+di, mono+di+tri and mono+di+tri+tetra-nucleotide compositions respectively. Moreover, j = 8, 9 correspond to features based on similarity between the query and reference strains (SF): j = 8 corresponds to genome similarity index dDDH and j = 9 corresponds to the index that relies on similarity between DNA gyrase (gyrB) genes. For each j-th GF, six TS+APS-based MSSPMs have been developed using six different ML algorithms. Because 9 sets of strains-based attributes (features) have been considered to describe the genome, 9 combined training sets (SQTS_cj_ j=1,9) have been used for model development. The j-th combined set SQTS_cj_ is created by uniting the j-th SQTS_i j_ set of eight MSs, that is, SQTS_cj_ =SQTS_1j_⋃ SQTS_2j_⋃…⋃ SQTS_8j_

### 2.7 Evaluation of the Quality of the Prediction

The following metrics have used to evaluate the quality of the prediction:

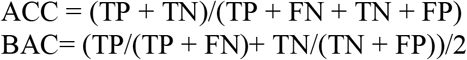

Where ACC is accuracy, BAC is balance accuracy, TP is true positive, TN is true negative, FP is false positive, and FN is true negative.

## 3. RESULTS

We have examined different ML algorithms for the development of MSS predictive models (MSSPM) for AMPs. The efficiency/precision of the model depends on the quality of the training set used in the learning process. We have followed two different approaches to create training sets for training MSSPMs. The first corresponds to choosing AMPs for which data on activity against certain microbial strains is available, and building training sets for them (sets of SQ type) by encoding sequences of the chosen AMPs. In case of strains with insufficient data, on the other hand, it is more reasonable to go the second way, which entails encoding information on microbial strains and adding it to the AMP sequence information to create samples for the training set. Instances that carry information on different pairs of TS+APS form training sets of SQTS type, where AMPs have been tested against the target strain from the pairs. SQ and SQTS training sets have been used to train various ML algorithms in order to reveal the most efficient combination “ML algorithm - type of encoding” (ML-TE).

### 3.1 Comparison of only AMP sequence-based MSSPMs

Six ML algorithms have been considered. Each trained on the eight training sets, created with the help of data on different MSs. Data on sequences of AMPs, tested on each MS has been used to create the eight training sets, SQi (i=1-8). The correspondence of i-th (SQi) training sets to particular MS and numbers of peptides in each learning (training) set are presented in the Table 1.

Each training set has been used to train each of the six different ML algorithms to reveal the most stable and accurate MSSPM. The quality of each MSSPM has been evaluated with the help of 10-fold cross validation. Traditional metrics have been used to assess MSSPMs performance. We define the models based solely on AMP sequences as belonging to the first group of MSSPMs and call them MSSPM_G1. Combination of eight training sets with six ML approaches gives 48 MSSPMs, and therefore MSSPM_G1 unites 48 models. Performance of each MSSPM is summarized in Table 2. One can see from this table that there is no single ML algorithm that works equally well (above 80% accuracy) for all training sets. Nevertheless, the predictive models developed using two ML algoritms — Random Forest and RealAdaBoost - show decent performance for majority of the datasets, with typical difference in accuracy between Random Forest-based and RealAdaBoost-based classifiers around 2 ± 1.3 %. For all strains except *Enterococcus faecalis ATCC 29212*, the accuracies of MSSPMs based on RealAdaBoost are higher than 80%.

**Table 2.** Accuracies attained by six ML algorithms, trained on SQi datasets(i=1,8)

| **Training set** | **RF\*** | **SVM\*** | **KN\*** | **RA\*** | **MP\*** | **DM\*** |
| --- | --- | --- | --- | --- | --- | --- |
| SQ<sub>1</sub> | 79.24 | 61.96 | 79.50 | 81.24 | 73.03 | 72.59 |
| SQ<sub>2</sub> | 82.24 | 69.64 | 77.78 | 83.41 | 77.01 | 73.03 |
| SQ<sub>3</sub> | 87.21 | 78.31 | 81.13 | 83.37 | 82.50 | 83.13 |
| SQ<sub>4</sub> | 81.48 | 67.55 | 84.36 | 84.05 | 81.28 | 77.02 |
| SQ<sub>5</sub> | 73.44 | 63.74 | 76.14 | 72.66 | 78.70 | 69.11 |
| SQ<sub>6</sub> | 80.64 | 64.16 | 78.02 | 82.24 | 75.05 | 71.52 |
| SQ<sub>7</sub> | 77.17 | 63.68 | 74.81 | 76.73 | 75.52 | 73.01 |
| SQ<sub>8</sub> | 81.98 | 54.97 | 79.48 | 81.68 | 79.64 | 81.06 |

\* RF -Random Forest; SVM – LibSVM; KN – K-nearest neighbours; RA – RealAdaBoost; MP – MultilayerPerceptron; DM – Dl4jMlpClassifier.

The models based on the LibSVM algorithm were quite inefficient for almost all datasets. It is perhaps worth noting, that the sets SQ_3_, SQ_4_, SQ_5_, SQ_7_, SQ_8_ are not very large and consequently our conclusions regarding efficiency of the models trained on them should be considered preliminary.

### 3.2 Comparison of TS+APS - based MSSPMs

Nine genome features (GF) have been used to encode microbial strain genomes and, together with AMP sequence encodings, create instances for TS +APS - based datasets. Seven of nine GF describe genome’s oligonucleotide composition and two give genome’s similarity with another genome. Each training set consists of data on all MS. Therefore, they are referred to as combined sets (SQTS_cj_ j=1,9). The datasets differ only by the encoding of MS genomes. Because 9 sets of MS-based features were considered, 9 combined datasets have been used to train the models. With six different machine learning algorithms, this makes 54 different models. To test the models’ performance, 10-fold cross validation has been performed. The given group of 54 MSSPMs will be referred to as MSSPMs G2. The accuracies of MSSPM G2 models are presented in Table 3, whose entries can be interpreted as prediction accuracies, averaged over different microbial strains. The accuracies for each particular strain can also be assessed using the corresponding SQTS_ij_ dataset as the test set. We have done so for a subset of MSSPMs G2 group, namely the sets of models G25, G28 and G29 each containing 6 different ML models, trained on the SQTS_c5_, SQTS_c8_ and SQTS_c9_ datasets respectively. The performance of these models has been assessed on the sets SQTS_ij_ with i=1,…,8, where the index i counts strains, and the index j corresponds to the given GF (j = 5, 8 and 9 for SQTS_c5_, SQTS_c8_ and SQTS_c9_ respectively). The total number of accuracy values computed is thus 18(the number of models) x 8(number of test sets for each model)), which sums up to 144 assessments. These values are presented in Fig. 1 and in the Table S2.

**Figure 1.**
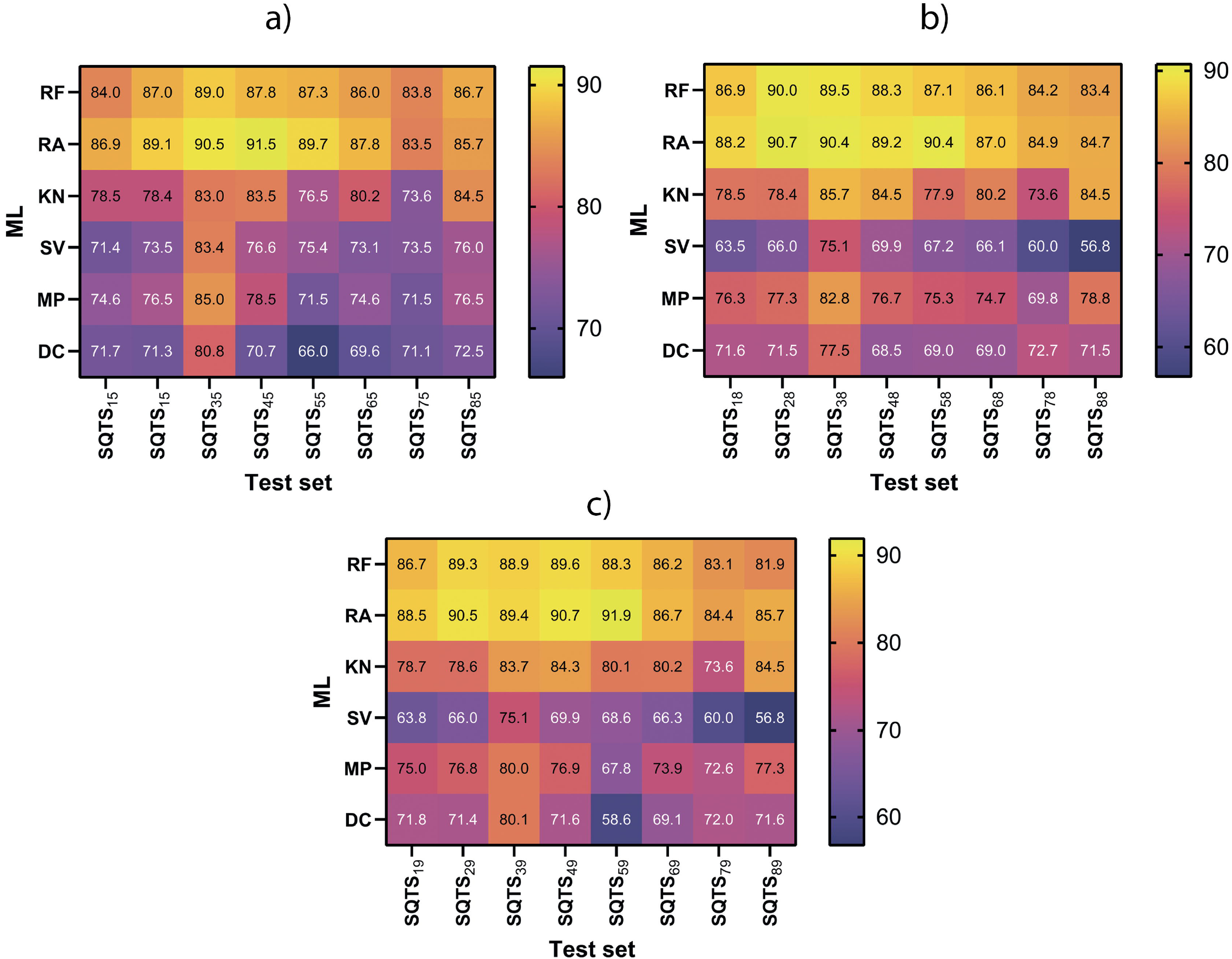
Balance accuracies for the models from three subgroups of the MSSPM G2 group: a) G25, b) G28 and c) G29 (abbreviations are described in Section 3.2). Test sets SQTSij were created based on data corresponding to a particular pair (i,j): i-th strain and j-th GF. The first index takes on the following values: i=1,…8, with i =1 corresponding to Escherichia coli ATCC 25922, i = 2 - Pseudomonas aeruginosa ATCC 27853, i = 3 - to Klebsiella pneumoniae ATCC 700603, i = 4 - to Salmonella typhimurium ATCC 14028, i = 5 - to Acinetobacter baumannii ATCC 19606, i = 6 - to Staphylococcus aureus ATCC 25923, i = 7 - to Enterococcus faecalis ATCC 29212, and i = 8 - to Bacillus subtilis ATCC 6633. The second index takes on the values j=5, 8, 9, where j = 5 corresponds to mono+di nucleotide compositions and j = 8, 9 correspond to SF: j = 8 corresponds to genome similarity index dDDH and j = 9 corresponds to index that relies on similarity between gyrB genes; SQTScj = SQTS1j ⋃ SQTS2j ⋃… ⋃ SQTS8j. BAC was evaluated using 10-fold cross-validation.

**Table 3.**
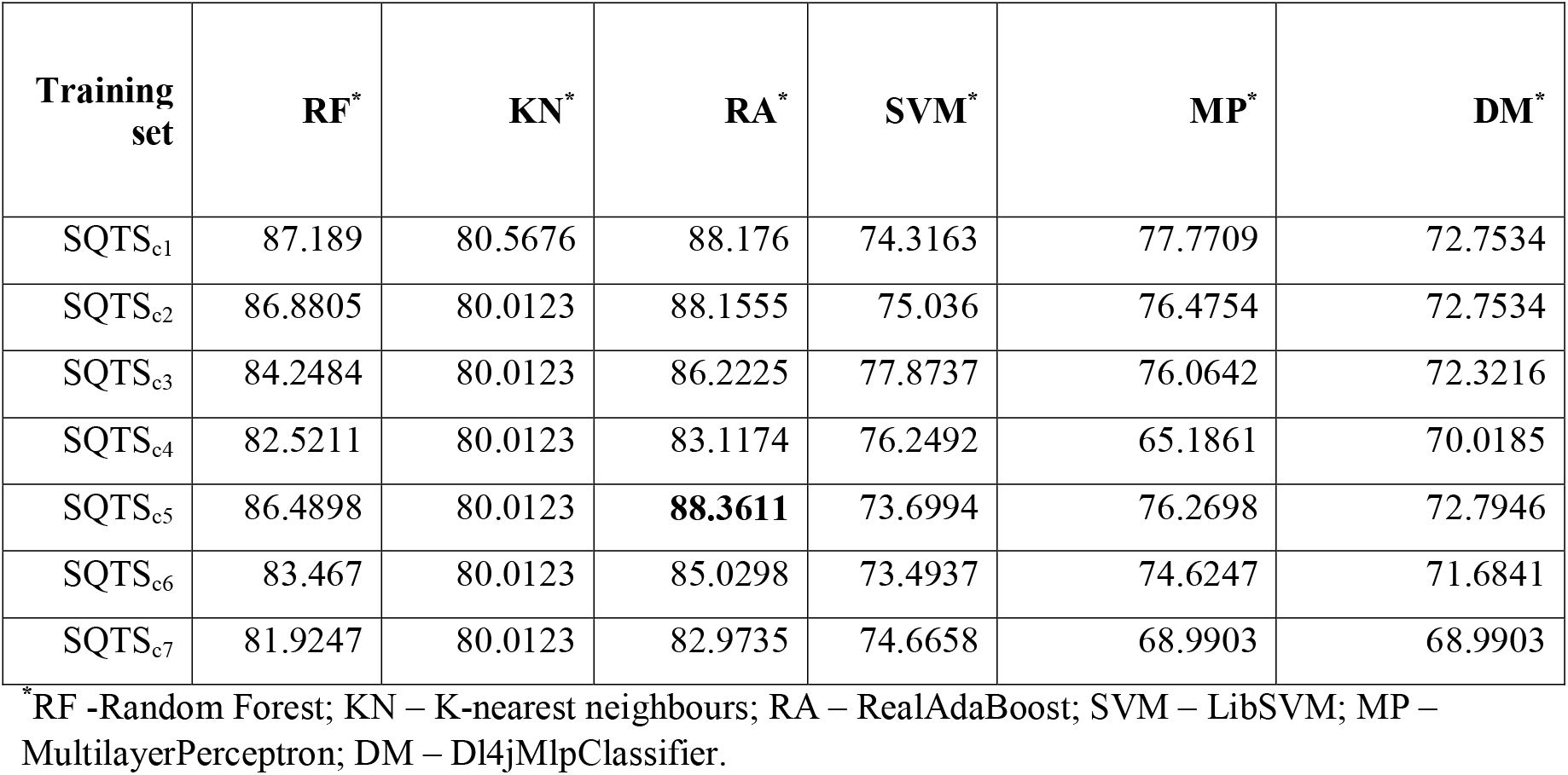
Performance of oligonucleotide composition-based predictive models based on 10-fold cross-validation (the largest value is highlighted in bold)

Taking into account the problem of insufficient data for many MS, we decided to develop other MSSPMs by training models on what we call ^i-^SQTS_cj_ training sets, where we have used the following definition: ^i-^SQTS_cj_ = SQTS_cj_ - SQTS_ij_. That is, we train the model by removing instances encoding a particular TS + APS pair (represented by the index i) and use these instances as the blind test set (repeating this procedure 8 times for each genome feature, represented by the index j). The corresponding group of MSSPM (6 ML algorithms, trained on 8×9 different datasets) is referred to as MSSPM G3 below. Similarly to the case of MSSPM G2 discussed above, the performance of the subset G35, G38 and G39 of these models has been assessed on the test sets ^i-^SQTS_cj_ with i=1,…,8. The results are presented in Table S3.

#### 3.2.1 Oligonucleotide composition-based predictive models

Seven kinds of oligonucleotide composition-based features have been considered to assess their impact on the performance of the six algorithms that form the base of our MSSPMs. The considered compositions are mono, di, tri and tetra-nucleotide compositions separately, as well as the following combinations of these compositions: (mono+di, mono+di+tri, mono+di+tri+ tetra). In Table 3, the performance metrics of the resulting 7 (training sets) * 6 (algorithms) = 42 models are presented. Again, all accuracies are derived from 10-fold cross-validation. It is clear from the results presented in Tables 2 and 3, that including additional information on genomes of microbial strains into the training sets improves performance of the corresponding MSSPM. Although these 42 models do not differ significantly from each other in terms of performance, the quality of models based on mono and di-nucleotide compositions was, on average, slightly better. Among the ML algorithms, Random Forest and RealAdaBoost demonstrated better performance, similar to our results with peptide-based features only.

Taking into account all of the above, we decided to use mono+di - nucleotide composition-based models to compare with inter-strain similarity-based predictive models.

#### 3.2.2 Inter-strain similarity-based predictive models

An alternative way to encode target strain genome is by using features that describe inter-strain similarity (SF). SF were represented as 5-dimensional vectors showing similarity between target genome and five (RGs).

The SQTS_c8_ and SQTS_c9_ sets have been created on the basis of encoding TS genomes with similarity features such as similarity indexes dDDH and gyrB gene similarity, assessed between TS and RG. Tables S1-S2 and Fig. 1–2 give accuracies and balance accuracies of predictive models based on various machine learning algorithms, trained on SQTS_c8_ and SQTS_c9_ datasets. In addition, the best model among oligo-nucleotide composition-based sets (trained on the SQTS_c5_) have been included as well.

Comparison of predictive models from MSSPM_G1 and MSSPM_G2 groups, built on two different encodings (AMP sequence-based and TS+APS-based respectively) demonstrated that addition of the TS genome-based attributes had positive effect on prediction accuracy (the accuracy increases by 3 to 17 percent depending on the precise strain with the increase of 5-7% for most strains, see Fig. 4). Balance accuracies for models from the MSSPM_G2 group vary from 84 to 92 % for different strains.

To see what the impact of missing data on a particular strain in a model’s training set is on the model’s performance, we can compare balance accuracies of the models from MSSPM_G2 and MSSPM_G3 (the latter group corresponds to models, trained on data with missing information about particular strains - see the definition above). The performance of models from these two groups are given in Fig. 1 and Fig. 2 respectively. To make comparison more reliable, only strains with abundant data were picked as test sets for the two sets of models (Table 1). Strains *Escherichia coli ATCC 25922, Pseudomonas aeruginosa ATCC 27853* and *Staphylococcus aureus ATCC 25923* were used for these purposes. The results show that balance accuracies of models from MSSPM_G3 on the given test sets (81-83%) are about 7% lower than balance accuracies of the models from MSSPM_G2 on the same test sets (88-90%). (We emphasise once again, that in the given procedure, models from the group MSSPM_G3 were tested on data that they haven’t ‘seen’ at their training time.) Therefore, we can conclude that our models’ prediction results should be quite reliable for (the majority of) new strains for which the data on antimicrobial activity is lacking.

**Figure 2.**
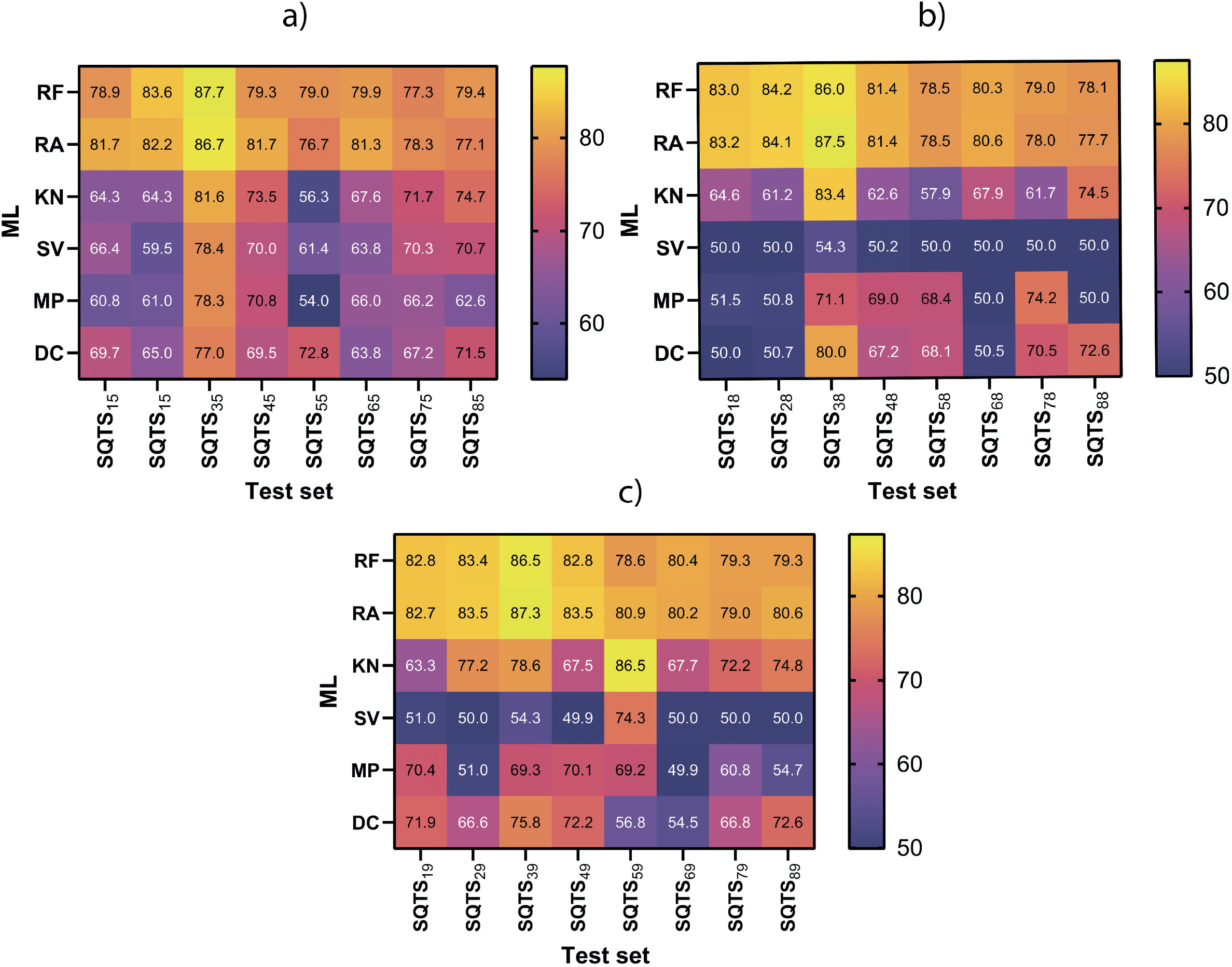
Balance accuracies for the models from three subgroups of the MSSPM G3 group: a) G35, b) G38 and c) G39 (abbreviations are described in Section 3.2). Test sets SQTSij were created based on data corresponding to a particular pair (i,j): i-th strain and j-th GF. The first index takes on the following values: i=1,…8, with i =1 corresponding to *Escherichia coli ATCC 25922*, i = 2 - *Pseudomonas aeruginosa ATCC 27853*, i = 3 - to *Klebsiella pneumoniae ATCC 700603*, i = 4 - to *Salmonella typhimurium ATCC 14028*, i = 5 - to *Acinetobacter baumannii ATCC 19606*, i = 6 - to *Staphylococcus aureus ATCC 25923*, i = 7 - to *Enterococcus faecalis ATCC 29212*, and i = 8 - to *Bacillus subtilis ATCC 6633*. The second index takes on the values j=5, 8, 9, where j = 5 corresponds to mono+di nucleotide compositions and j = 8, 9 correspond to SF: j = 8 corresponds to genome similarity index dDDH and j = 9 corresponds to index that relies on similarity between gyrB genes. BAC was evaluated using 10-fold cross-validation.

We should perhaps note that there is no single genome-based set of features that lead to significantly better performance compared to the others. As to the ML algorithms, Random Forest and RealAdaBoost still give the best prediction results, just as they did for the case of purely sequence-based predictive models.

### 3.3 Comparison of MSSPMs with other available tools

We have surveyed available tools that predict AMPs against particular microbial strains or species and compared them to the best models developed in the current work. Four predictive tools [5,12–14] have been selected. The algorithm from ref. [12] provides a score of the effectiveness of a peptide sequence against various microbial species. However, from this score it is not clear if the peptide is active against particular strain or not and so we did not use this tool for comparison. Three remaining algorithms [5,13–14] perform prediction of anti-tubercular peptides. Among these, only two offer on-line prediction capability [12,14]. Akbar et al., mentioned [5] that the source code for their model is publicly available in a github repository, but the corresponding link appears to be broken. Therefore, we decided to compare our best models — those based on the RF and RealAdaBoost algorithms and trained on the SQTS_c5_, SQTS_c8_ and SQTS_c9_ datasets — with the models presented as anti-tubercular peptide prediction tools: AntiTB [13] and Atb [14]. AntiTB server offer four models for prediction (**AntiTB_MD SVM ensemble, AntiTB_RD SVM ensemble, AntiTB_MD Hybrid and AntiTB_RD Hybrid**) that differ by the corresponding training sets and ML algorithms. Atb server, on the other hand, offers two models (**Atb_MD and Atb_RD**) that have been trained on what is referred to as ‘MD’ and ‘MR’ datasets. We have to note, that the MD and MR training sets have been shared between models from both of the above-mentioned tools (AntiTB and Atb). MD and MR have a shared positive set, created on the basis of data from the AntiTB database [41], while the negative sets are different, created on the basis of SWISS-PROT [42] and DBAASP [24] respectively. It is worth noting that negative sets consist of peptides not tested against TB.

To perform the comparison, a benchmark test set (BTS) has to be chosen. According to our definition, laid out in the method section, a set of anti-TB peptides, forming the positive set for BTS must consist of AMPs tested on Mycobacteria TB (M.tb) and having MIC < 25 μg/mL. The data on separate strains of M.tb is scarce, therefore we have used data on two strains of M.tb from DBAASP, namely H37v and lux, and peptides that have MIC < 25 against one of these strains has been involved in the positive set of BTS. Negative set consists of peptides that have been tested on H37v and lux strains and that have MIC > 100 for both. Consequently, we have created a benchmark test set, which consist of 74 peptides: 38 active peptides for the positive set and 36 non-active peptides for the negative set. M.tb -specific predictive models of AMP, together with other models developed and described in this work have been tested on the given BTS (Table 4). It is worth noting that the positive set from the BTS is a truly blind test set for our MSSPMs (because our MSSPMs have not seen this data at training time), while it is not for anti-tubercular peptide predictive models AntiTB and Atb.

**Table 4.** Performances of different predictive models on the MT-DS set for Benchmark test

| Predictive models | Sensitivity (%) | Specificity (%) | Balance Accuracy (%) | Accuracy (%) | Ref: |
| --- | --- | --- | --- | --- | --- |
| AntiTB_MD SVM ensemble* | 97.37 | 38.89 | 68.13 | 68.92 | [13] |
| AntiTB_RD SVM ensemble* | 97.37 | 47.22 | 72.30 | 72.97 | [13] |
| AntiTB_MD Hybrid method* | 97.37 | 36.11 | 66.74 | 67.57 | [13] |
| AntiTB_RD Hybrid method* | 100.00 | 30.56 | 65.28 | 66.22 | [13] |
| Atb_MD* | 97.37 | 47.22 | 72.30 | 72.97 | [14] |
| Atb_RD* | 97.37 | 25.00 | 61.18 | 62.16 | [14] |
| Random Forest on SQTS<sub>c5</sub> | 89.47 | 86.11 | 87.79 | 87.84 | This work |
| RealAdaBoost on SQTS<sub>c5</sub> | 92.11 | 86.11 | 89.11 | 89.19 | This work |
| Random Forest on SQTS<sub>c8</sub> | 94.74 | 88.89 | 91.81 | 91.89 | This work |
| RealAdaBoost on SQTS<sub>c8</sub> | 92.11 | 88.89 | 90.50 | 90.54 | This work |
| Random Forest on SQTS<sub>c9</sub> | 92.11 | 88.89 | 90.50 | 90.54 | This work |
| RealAdaBoost on SQTS<sub>c9</sub> | 92.11 | 88.89 | 90.50 | 90.54 | This work |

\* **AntiTB** – server from [13], **Atb** – server from [14], **MD** – data for the negative set from DBAASP, **RD** – data for the negative set from SWISS-PROT

The test results show that anti-tubercular AMP prediction servers’ performance in the prediction of the active AMP (i.e. sensitivity) is higher than our models’ performance. However, AMP models underperform when non-anti-M.tb peptide is to be predicted. Consequently, the specificities and accuracies of the anti-M.tb AMP prediction tools are markedly lower than the same characteristics of our models.

These results can be explained as follows. Learning on the positive training sets allows anti-tubercular AMP prediction methods to learn shared features of AMPs. These shared features are thought to be linked with the capability to be active against an “averaged” membrane [7]. Therefore, the trained model predicts all membrane-active peptides as AMP, although it is not correct, because in reality not all membrane-active peptides are AMPs. If the model doesn’t know these additional important properties, it can’t separate membrane-active non-AMPs from AMPs. When randomly selected ‘putative non-AMP sequences’ (that is, reasonably supposed to be non-membrane active) have been used as a negative training set, the developed models distinguish membrane-active peptides from non-membrane-active ones, with high efficiency. But the same model fails when it is necessary to distinguish non-active against a particular strain but membrane-active peptide from AMP active against the same strain.

As a disclaimer, the definition of a ‘putative non-AMP sequence’ might be unavoidably incomplete: even if it can be proven that a certain AMP is not active against *n* different microbes, it can still turn out to be active against some (n+1)-th microbe, which has not been yet tested. Consequently, for a model to be capable of distinguishing non-active against a particular strain but membrane-active peptide from AMP, it has to be trained on a set that contains non-AMP (including membrane-active ones), tested on a particular strain.

We have to emphasise, that natural AMPs evolve in a certain habitat, aiming to be active against a particular microbial set. At the same time, they might not be active against others. Moreover, synthetic non-AMPs were designed with an aim for them to be active against a particular set of microbes, but without success. Therefore, many peptides from databases, that were assessed on antimicrobial potency and turned out not active carry some features shared with AMP and consequently with membrane-active peptides. Therefore, training on the negative training set that consists of experimentally proven non-AMPs will give the model knowledge on the features of membrane-active but non-AMP peptides that distinguish them from AMP.

This is why all models trained on “putative non-AMP” sets without specifying the target, that is the particular strain (type of membrane) they have to interact with, fail when non-AMP peptides have to be predicted. Many peptides can be active against a biological membrane in general, but only particular ones can have appropriate effect on a particular membrane.

## 4. CONCLUSION

To overcome the problem of drug-resistance, new paradigms and approaches are needed. One approach is to learn the lessons from AMPs, a class of natural molecules that have the experience to combat pathogenic microbes without known risk of the development of resistance. Computational approaches are widely used to aid and complement the effort to develop new peptide-based antibiotics. Recent innovations in ML algorithms and the availability of better quality AMP datasets have caused proliferation of ML-aided prediction methods that aim to perform rational design of antibiotics possessing new mechanisms of action. Xu et all [1] in their work survey more than 30 approaches for AMP identification and try to evaluate the predictive performance of the tools based on an independent test dataset. Predictive performance of the majority of methods on the test sets used is high, but some drawbacks of the approaches make them less applicable to target-oriented design of AMP. The major drawback is that all predictive models surveyed in [1] have been trained in non-microbial-strain-specific training sets. This is a problem, as the potency of peptides (widely presented as MIC), and consequently their class, is exclusively determined on the microbial strain level. Therefore, positive training sets consisting of AMPs, active against some microbial strains but not active against others will fail to provide comprehensive learning (essentially because AMPs carry uncertainty in the definition of the class of peptide). Even more problematic is creation of a statistically reliable, experimentally proven negative training sets. It is reasonable to suppose that patterns, learned on the non-microbial-strain-specific training sets are only sensitive to shared features of the AMP, for instance features linked with the activity against “average” membrane, but knowledge on the full set of features necessary to perform antimicrobial action against certain strain (i.e. particular membrane) is absent.

Development of predictive models, based on the microbial strain-specific training sets is complicated due to insufficient data on the majority of microbial strains. To overcome the problem of insufficiency of data, we have developed microbial strain-specific predictive models relying on the attributes characterizing both the AMP sequences and target microbial-strain genomes at the same time. Prediction accuracies of models trained on datasets involving TS genome-based attributes have risen on average by 7% with respect to models relying on AMP sequence information only. Among ML approaches, Random Forest and RealAdaBoost give the best prediction results. Our models allow to perform prediction of AMPs against microbial strain with insufficient data.

The current performance of the models can be improved in multiple ways: i) by expanding the volume of data for the training set ii) by increasing the number of reference genomes and iii) by developing a new encoding system aiming to describe AMP sequence and TS genome more comprehensively. We invite researchers to use our on-line prediction tool and plan to continue upgrading it as outlined above.

## DATA AVAILABILITY

The MSS AMP predictors that correspond to the most accurate and stable models are available on DBAASP website: https://dbaasp.org/tools?page=genome-prediction

## ACKNOWLEDGEMENTS

This work was supported with funds from the National Institute of Allergy and Infectious Diseases (NIAID), National Institutes of Health, Department of Health and Human Services and the International Science and Technology Center (G-2102).

## REFERENCES

1. Xu J, Li F, Leier A, Xiang D, et al. Comprehensive assessment of machine learningbased methods for predicting antimicrobial peptides. Briefings in Bioinformatics 2021;22:bbab083, https://doi.org/10.1093/bib/bbab083

2. Pinacho-Castellanos SA, García-Jacas CR, Gilson MK, et al. Alignment-Free Antimicrobial Peptide Predictors: Improving Performance by a Thorough Analysis of the Largest Available Data Set. Journal of Chemical Information and Modeling 2021;61:3141–3157

3. Waghu FH, Barai RS, Gurung P, et al. CAMPR3: a database on sequences, structures and signatures of antimicrobial peptides. Nucleic Acids Res 2015;44: D1094–1097.

4. Kavousi K, Bagheri M, Behrouzi S, et al. IAMPE: NMR Assisted Computational Prediction of Antimicrobial Peptides. Journal of Chemical Information and Modeling 2020;60:4691–4701

5. Akbar S, Ahmada A Hayat M, et al. iAtbP-Hyb-EnC: Prediction of Antitubercular peptides Via Heterogeneous Feature Representation and Genetic Algorithm based Ensemble Learning Model. Computers in Biology and Medicine 2021;137:104778.

6. Guo Y, Yan K, Liu B. PreTP-EL: prediction of therapeutic peptides based on ensemble learning, Briefings in Bioinformatics 2021;bbab358, https://doi.org/10.1093/bib/bbab358

7. Lee EY, Fulan BM, Wong GC, et al. Mapping membrane activity in undiscovered peptide sequence space using machine learning. Proc Natl Acad Sci U S A 2016; 113:13588–13593. doi: 10.1073/pnas.1609893113.

8. Yan J, Bhadra P, Li A. et al. Deep-AmPEP30: Improve Short Antimicrobial Peptides Prediction with Deep Learning. Molecular Therapy - Nucleic Acids 2020;20:882–894. doi: 10.1016/j.omtn.2020.05.006.

9. Speck-Planche A, Kleandrova VV, Ruso JM, et al. First Multitarget Chemo-Bioinformatic Model To Enable the Discovery of Antibacterial Peptides against Multiple Gram-PositivePathogens J. Chem. Inf. Model 2016;56:588–598.

10. Kleandrova VV, Ruso JM, Speck-Planche A, et al. Enabling the Discovery and Virtual Screening of Potent and Safe Antimicrobial Peptides. Simultaneous Prediction of Antibacterial Activity and Cytotoxicity *ACS Comb*. Sci 2016;18:490–498.

11. Nantasenamat LHC. Toward insights on determining factors for high activity in antimicrobial peptides via machine learning. PeerJ 2019;7:e8265. https://doi.org/10.7717/peerj.8265

12. Gull S, Minhas FUAA. AMP0: Species-Specific Prediction of Anti-microbial Peptides using Zero and Few Shot Learning *IEEE/ACM Transactions on Computational Biology and Bioinformatics*, doi: 10.1109/TCBB.2020.2999399.

13. Usmani SS, Bhalla S, Raghava GPS. Prediction of Antitubercular Peptides From Sequence Information Using Ensemble Classifier and Hybrid Features. Front Pharmacol. 2018;9:954. doi:10.3389/fphar.2018.00954

14. Manavalan B, Basith S, Shin TH, et al. AtbPpred: A Robust Sequence-Based Prediction of Anti-Tubercular Peptides Using Extremely Randomized Trees. Comput Struct Biotechnol J. 2019;17:972–981. doi: 10.1016/j.csbj.2019.06.024.

15. Losin L, Veltri D. Exploring target specificity of antimicrobial peptides through deep learning embeddings. In: Proceedings of the 12th ACM Conference on Bioinformatics, Computational Biology, and Health Informatics. August 2021 Article No.: 77 P.1 doi: 10.1145/3459930.3469506.

16. Wang, C.; Garlick, S.; Zloh, M. Deep Learning for Novel Ant/imicrobial Peptide Design. Biomolecules 2021;11:471. https://doi.org/10.3390/biom11030471

17. Vishnepolsky B, Gabrielian A, Rosenthal A, et al. Predictive Model of Linear Antimicrobial Peptides Active against Gram-Negative Bacteria. Journal of Chemical Information and Modeling 2018;58:1141–1151 doi: 10.1021/acs.jcim.8b00118.

18. Vishnepolsky B, Grigolava M, Zaalishvili G, et al. DBAASP Special prediction as a tool for the prediction of antimicrobial potency against particular target species. in Proceedings of the 4th International Electronic Conference on Medicinal Chemistry, 1-30 November 2018, MDPI: Basel, Switzerland, doi:10.3390/ecmc-4-05608.

19. Weisburg WG, Barns SM, Pelletier DA, et al. 16S ribosomal DNA amplification for phylogenetic study. J Bacteriol. 1991;173:697–703. doi:10.1128/jb.173.2.697-703.1991

20. Akhurst RJ, Boemare NE, Janssen PH Peel, et al. Taxonomy of Australian clinical isolates of the genus Photorhabdus and proposal of Photorhabdus asymbiotica subsp. asymbiotica subsp. nov. and P. asymbiotica subsp. australis subsp. nov. International Journal of Systematic and Evolutionary Microbiology 2004;54:1301–1310.

21. Chun J, Rainey FA. Integrating genomics into the taxonomy and systematics of the Bacteria and Archaea. Int J Syst Evol Microbiol 2014;64:316–324.

22. Konstantinidis K, Tiedje J. Genomic insights that advance the species definition for prokaryotes. Proc. Natl. Acad. Sci. USA 2005;102:2567–2572.

23. Meier-Kolthoff JP, Auch AF, Klenk HP, et al. Genome sequence-based species delimitation with confidence intervals and improved distance functions. BMC Bioinformatics 2013;14:60

24. Pirtskhalava M, Amstrong AA, Grigolava M, et al. DBAASP v3: database of antimicrobial/cytotoxic activity and structure of peptides as a resource for development of new therapeutics. Nucleic Acids Research 2021;49:D288–D297

25. Benson DA, Cavanaugh M, Clark K, et al. GenBank Nucleic Acids Research 2013; 41:D36–D42

26. https://www.scientificamerican.com/article/who-releases-list-of-worlds-most-dangerous-superbugs/

27. Mulani MS, Kamble EE, Kumkar SN, et al. Emerging strategies to combat ESKAPE pathogens in the era of antimicrobial resistance: *A review*. Front. Microbiol. 2019;10:539.

28. Karlin, S. Global dinucleotide signatures and analysis of genomic heterogeneity. Current opinion in microbiology 1998;1:598–610.

29. Nakashima H, Nakashima K, Ooi T, et al. Diferences in Dinucleotide Frequencies of Human, Yeast, and Escherichia coli Genes. DNA Research 1997:4:185–192.

30. Nakashima H, Ota M, Nakashima K, et al. Genes from nine genomes are separated into their organisms in the dinucleotide composition space. DNA Research 1998;5:251–259.

31. Abe T, Kanaya S, Kinouchi M, et al. Informatics for unveiling hidden genome signatures. Genome Res 2003;13(4):693–702. doi:10.1101/gr.634603

32. Pride DT, Meinersmann RJ, Wassenaar TM, et al. Evolutionary implications of microbial genome tetranucleotide frequency biases. Genome Res 2003;13(2):145–158. doi:10.1101/gr.335003

33. Takahashi M, Kryukov K, Saitoua N, et al. Estimation of bacterial species phylogeny through oligonucleotide frequency distances. Genomics 2009;93:525–533.

34. http://ggdc.dsmz.de/ggdc_background.php#entry1

35. Fu LM, Fu-Liu CS. Is Mycobacterium tuberculosis a closer relative to Gram-positive or Gram-negative bacterial pathogens?. Tuberculosis 2002:82:85–90.

36. Wang G, Li X, Wang Z. APD3: the antimicrobial peptide database as a tool for research and education. Nucleic Acids Research 2016;44:D1087–D1093.

37. Jepson AK, Schwarz-Linek J, Ryan L, et al. What Is the ‘Minimum Inhibitory Concentration’ (MIC) of Pexiganan Acting on Escherichia coli?-A Cautionary Case Study. Adv Exp Med Biol 2016;915:33–48. doi: 10.1007/978-3-319-32189-9_4

38. Vishnepolsky B, Pirtskhalava M. Prediction of linear cationic antimicrobial peptides based on characteristics responsible for their interaction with the membranes. J. Chem. Inf. Model. 2014; 54:1512–1523.

39. Pirtskhalava M Vishnepolsky B, Grigolava M, et al. Physicochemical Features and Peculiarities of Interaction of AMP with the Membrane. Pharmaceuticals 2021;14:471.

40. Kyte J, Doolittle RF. A simple method for displaying the hydropathic character of a protein. J. Mol. Biol. 1982;157:105–132.

41. Usmani SS, Kumar R, Kumar V, et al. AntiTbPdb: a knowledgebase of anti-tubercular peptides. Database 2018:2018, bay025, https://doi.org/10.1093/database/bay025

42. The UniProt Consortium. UniProt: the universal protein knowledgebase in 2021. Nucleic Acids Res. 2021;49:D1

