## Supplemental Tables for "Comparative analysis of machine learning algorithms on the microbial strain-specific AMP prediction"

*Corresponding Authors:

Table S1. The number of the peptides with data of antimicrobial activities for the selected microbial strains.

| **Selected strains** | **Number of the peptides** |
|---|---|
| Escherichia coli ATCC 25922 | 5,563 |
| Staphylococcus aureus ATCC 25923 | 3,582 |
| Pseudomonas aeruginosa ATCC 27853 | 3,691 |
| Klebsiella pneumoniae ATCC 700603 | 817 |
| Salmonella typhimurium ATCC 14028 | 586 |
| Enterococcus faecalis ATCC 29212 | 1,038 |
| Acinetobacter baumannii ATCC 19606 | 940 |
| Bacillus subtilis ATCC 6633 | 729 |

Table S2. Prediction performances (Balance accuracies and accuracies) of the models of three subugroups of MSSPM G2 group (G25, G28, G29).

| | **RF** | | **RA** | | **KN** | | **SVM** | | **MP** | | **DM** | |
|---|---|---|---|---|---|---|---|---|---|---|---|---|
| | **BAC** | **ACC** | **BAC** | **ACC** | **BAC** | **ACC** | **BAC** | **ACC** | **BAC** | **ACC** | **BAC** | **ACC** |
| $SQTS_{15}$ | 84.0 | 84.8 | 86.9 | 87.4 | 78.5 | 79.3 | 71.4 | 71.6 | 74.6 | 75.4 | 71.7 | 73.1 |
| $SQTS_{15}$ | 87.0 | 87.0 | 89.1 | 89.1 | 78.4 | 78.4 | 73.5 | 73.5 | 76.5 | 76.5 | 71.3 | 71.3 |
| $SQTS_{35}$ | 89.0 | 88.4 | 90.5 | 90.2 | 83.0 | 83.2 | 83.4 | 83.8 | 85.0 | 85.0 | 80.8 | 80.3 |
| $SQTS_{45}$ | 87.8 | 88.7 | 91.5 | 92.0 | 83.5 | 84.4 | 76.6 | 78.2 | 78.5 | 79.3 | 70.7 | 72.7 |
| $SQTS_{55}$ | 87.3 | 92.5 | 89.7 | 93.1 | 76.5 | 83.9 | 75.4 | 73.0 | 71.5 | 76.4 | 66.0 | 78.7 |
| $SQTS_{65}$ | 86.0 | 86.7 | 87.8 | 88.4 | 80.2 | 80.7 | 73.1 | 73.3 | 74.6 | 75.3 | 69.6 | 70.5 |
| $SQTS_{75}$ | 83.8 | 84.5 | 83.5 | 84.2 | 73.6 | 74.0 | 73.5 | 74.6 | 71.5 | 71.6 | 71.1 | 71.9 |
| $SQTS_{85}$ | 86.7 | 88.2 | 85.7 | 87.8 | 84.5 | 86.1 | 76.0 | 76.3 | 76.5 | 81.6 | 72.5 | 79.6 |

| | | | | | | | | | | | | |
|---|---|---|---|---|---|---|---|---|---|---|---|---|
| SQTS$_{c5}$ | 85.9 | 86.5 | 88.0 | 88.4 | 79.5 | 80.0 | 73.6 | 73.7 | 75.7 | 76.3 | 71.8 | 72.8 |
| SQTS$_{18}$ | 86.9 | 87.1 | 88.2 | 88.5 | 78.5 | 79.3 | 63.5 | 67.2 | 76.3 | 77.0 | 71.6 | 73.1 |
| SQTS$_{28}$ | 90.0 | 90.0 | 90.7 | 90.7 | 78.4 | 78.4 | 66.0 | 66.1 | 77.3 | 77.3 | 71.5 | 71.5 |
| SQTS$_{38}$ | 89.5 | 89.0 | 90.4 | 90.2 | 85.7 | 85.5 | 75.1 | 72.8 | 82.8 | 82.7 | 77.5 | 78.0 |
| SQTS$_{48}$ | 88.3 | 89.5 | 89.2 | 90.2 | 84.5 | 85.1 | 69.9 | 73.1 | 76.7 | 78.5 | 68.5 | 70.9 |
| SQTS$_{58}$ | 87.1 | 90.2 | 90.4 | 93.1 | 77.9 | 83.9 | 67.2 | 80.5 | 75.3 | 77.0 | 69.0 | 75.9 |
| SQTS$_{68}$ | 86.1 | 86.7 | 87.0 | 87.5 | 80.2 | 80.7 | 66.1 | 69.3 | 74.7 | 75.3 | 69.0 | 70.1 |
| SQTS$_{78}$ | 84.2 | 85.1 | 84.9 | 85.7 | 73.6 | 74.0 | 60.0 | 62.4 | 69.8 | 70.1 | 72.7 | 73.1 |
| SQTS$_{88}$ | 83.4 | 86.1 | 84.7 | 87.3 | 84.5 | 86.1 | 56.8 | 71.4 | 78.8 | 83.3 | 71.5 | 79.2 |
| SQTS$_{c8}$ | 87.3 | 87.7 | 88.4 | 88.8 | 79.6 | 80.1 | 65.4 | 68.4 | 76.2 | 76.8 | 71.5 | 72.5 |
| SQTS$_{19}$ | 86.7 | 86.9 | 88.5 | 88.7 | 78.7 | 79.6 | 63.8 | 67.5 | 75.0 | 76.0 | 71.8 | 73.0 |
| SQTS$_{29}$ | 89.3 | 89.3 | 90.5 | 90.5 | 78.6 | 78.6 | 66.0 | 66.1 | 76.8 | 76.8 | 71.4 | 71.4 |
| SQTS$_{39}$ | 88.9 | 88.4 | 89.4 | 89.0 | 83.7 | 83.8 | 75.1 | 72.8 | 80.0 | 79.8 | 80.1 | 80.3 |
| SQTS$_{49}$ | 89.6 | 90.5 | 90.7 | 91.3 | 84.3 | 84.7 | 69.9 | 73.1 | 76.9 | 77.8 | 71.6 | 72.7 |
| SQTS$_{59}$ | 88.3 | 92.0 | 91.9 | 94.3 | 80.1 | 85.1 | 68.6 | 80.5 | 67.8 | 79.3 | 58.6 | 77.0 |
| SQTS$_{69}$ | 86.2 | 86.9 | 86.7 | 87.3 | 80.2 | 80.7 | 66.3 | 69.4 | 73.9 | 74.7 | 69.1 | 70.1 |
| SQTS$_{79}$ | 83.1 | 84.2 | 84.4 | 85.4 | 73.6 | 74.0 | 60.0 | 62.4 | 72.6 | 73.1 | 72.0 | 72.8 |
| SQTS$_{89}$ | 81.9 | 84.5 | 85.7 | 87.8 | 84.5 | 86.1 | 56.8 | 71.4 | 77.3 | 81.6 | 71.6 | 78.8 |
| SQTS$_{c9}$ | 87.0 | 87.6 | 88.4 | 88.8 | 79.7 | 80.2 | 65.5 | 68.5 | 75.6 | 76.3 | 71.7 | 72.6 |

Test sets SQTSij were created by the data of particular pair of (i,j) (i-th organism and j-th GF),where i=1,…8, and i=1 correspond to *Escherichia coli ATCC 25922*, 2 – *Pseudomonas aeruginosa ATCC 27853*, 3 – to *Klebsiella pneumoniae ATCC 700603*, 4 – to *Salmonella typhimurium ATCC 14028*, 5 – to *Acinetobacter baumannii ATCC 19606*, 6 – to *Staphylococcus aureus ATCC 25923*, 7 – to *Enterococcus faecalis ATCC 29212*, 8 – to *Bacillus subtilis ATCC 6633*, respectively; j=5, 8, 9, where j=5- corresponds to mono+di nucleotide compositions and j=8,9 corresponds to SF: 8 – corresponds to genome similarity index dDDH and 9 – corresponds to index relied on similarity between DNA gyrase genes; $SQTS_{cj} = SQTS_{1j} \cup SQTS_{2j} \cup \ldots \cup SQTS_{8j}$

RF -Random Forest; SVM – LibSVM; KN – K-nearest neighbours; RA – RealAdaBoost; MP – MultilayerPerceptron; DM – Dl4jMlpClassifier

BAC and ACC was evaluated using 10-fold cross-validation.

Table S3. Prediction performances (Balance accuracies and accuracies) of the models of three subugroups of MSSPM G3 group (MSSPM G35, MSSPM G38 MSSPM G39).

| | **RF** | | **RA** | | **KN** | | **SVM** | | **MP** | | **DM** | |
|---|---|---|---|---|---|---|---|---|---|---|---|---|
| | **BAC** | **ACC** | **BAC** | **ACC** | **BAC** | **ACC** | **BAC** | **ACC** | **BAC** | **ACC** | **BAC** | **ACC** |
| SQTS$_{15}$ | 78.9 | 79.5 | 81.7 | 81.9 | 64.3 | 63.1 | 66.4 | 64.4 | 60.8 | 65.0 | 69.7 | 71.4 |
| SQTS$_{25}$ | 83.6 | 83.6 | 82.2 | 82.2 | 64.3 | 64.2 | 59.5 | 59.4 | 61.0 | 60.9 | 65.0 | 64.9 |
| SQTS$_{35}$ | 87.7 | 87.3 | 86.7 | 86.1 | 81.6 | 82.7 | 78.4 | 79.8 | 78.3 | 76.9 | 77.0 | 76.3 |
| SQTS$_{45}$ | 79.3 | 80.0 | 81.7 | 82.5 | 73.5 | 75.6 | 70.0 | 71.6 | 70.8 | 69.1 | 69.5 | 71.3 |
| SQTS$_{55}$ | 79.0 | 84.5 | 76.7 | 81.0 | 56.3 | 55.2 | 61.4 | 58.6 | 54.0 | 51.7 | 72.8 | 71.3 |
| SQTS$_{65}$ | 79.9 | 80.2 | 81.3 | 81.4 | 67.6 | 66.8 | 63.8 | 61.8 | 66.0 | 65.5 | 63.8 | 59.9 |
| SQTS$_{75}$ | 77.3 | 79.1 | 78.3 | 80.0 | 71.7 | 73.7 | 70.3 | 71.9 | 66.2 | 67.2 | 67.2 | 69.6 |
| SQTS$_{85}$ | 79.4 | 79.6 | 77.1 | 78.0 | 74.7 | 72.7 | 70.7 | 66.1 | 62.6 | 60.4 | 71.5 | 61.6 |

| | | | | | | | | | | | | |
|---|---|---|---|---|---|---|---|---|---|---|---|---|
| $SQTS_{18}$ | 83.0 | 82.5 | 83.2 | 82.8 | 64.6 | 65.8 | 50.0 | 58.1 | 51.5 | 59.1 | 50.0 | 58.1 |
| $SQTS_{28}$ | 84.2 | 84.3 | 84.1 | 84.2 | 61.2 | 61.3 | 50.0 | 50.3 | 50.8 | 51.1 | 50.7 | 51.0 |
| $SQTS_{38}$ | 86.0 | 85.0 | 87.5 | 86.7 | 83.4 | 82.1 | 54.3 | 48.6 | 71.1 | 72.3 | 80.0 | 79.8 |
| $SQTS_{48}$ | 81.4 | 82.2 | 81.4 | 82.2 | 62.6 | 62.9 | 50.2 | 54.9 | 69.0 | 71.6 | 67.2 | 69.8 |
| $SQTS_{58}$ | 78.5 | 82.8 | 78.5 | 82.8 | 57.9 | 58.6 | 50.0 | 26.4 | 68.4 | 67.8 | 68.1 | 73.6 |
| $SQTS_{68}$ | 80.3 | 80.4 | 80.6 | 80.6 | 67.9 | 67.0 | 50.0 | 56.5 | 50.0 | 56.5 | 50.5 | 44.1 |
| $SQTS_{78}$ | 79.0 | 80.6 | 78.0 | 79.7 | 61.7 | 61.5 | 50.0 | 54.6 | 74.2 | 73.4 | 70.5 | 71.9 |
| $SQTS_{88}$ | 78.1 | 78.8 | 77.7 | 78.8 | 74.5 | 72.2 | 50.0 | 31.0 | 50.0 | 69.0 | 72.6 | 63.3 |
| $SQTS_{19}$ | 82.8 | 82.2 | 82.7 | 82.3 | 63.3 | 65.0 | 51.0 | 58.5 | 70.4 | 69.3 | 71.9 | 72.1 |
| $SQTS_{29}$ | 83.4 | 83.4 | 83.5 | 83.5 | 77.2 | 77.3 | 50.0 | 50.3 | 51.0 | 51.3 | 66.6 | 66.7 |
| $SQTS_{39}$ | 86.5 | 85.5 | 87.3 | 86.7 | 78.6 | 78.6 | 54.3 | 48.6 | 69.3 | 70.5 | 75.8 | 74.6 |
| $SQTS_{49}$ | 82.8 | 83.3 | 83.5 | 84.0 | 67.5 | 66.9 | 49.9 | 54.5 | 70.1 | 71.3 | 72.2 | 72.7 |
| $SQTS_{59}$ | 78.6 | 83.9 | 80.9 | 86.2 | 86.5 | 87.4 | 74.3 | 68.4 | 69.2 | 79.3 | 56.8 | 76.4 |
| $SQTS_{69}$ | 80.4 | 80.5 | 80.2 | 79.9 | 67.7 | 66.9 | 50.0 | 56.5 | 49.9 | 43.4 | 54.5 | 48.7 |
| $SQTS_{79}$ | 79.3 | 80.9 | 79.0 | 80.6 | 72.2 | 74.3 | 50.0 | 54.6 | 60.8 | 63.0 | 66.8 | 69.3 |
| $SQTS_{89}$ | 79.3 | 80.0 | 80.6 | 81.2 | 74.8 | 72.2 | 50.0 | 31.0 | 54.7 | 38.0 | 72.6 | 63.3 |

Test sets SQTSij were created by the data of particular pair of (i,j) (i-th organism and j-th GF),where i=1,…8, and i=1 correspond to Escherichia coli ATCC 25922, 2 – Pseudomonas aeruginosa ATCC 27853, 3 – to Klebsiella pneumoniae ATCC 700603, 4 – to Salmonella typhimurium ATCC 14028, 5 – to Acinetobacter baumannii ATCC 1960, 6 – to Staphylococcus aureus ATCC 25923, 7 – to Enterococcus faecalis ATCC 29212, 8 – to Bacillus subtilis ATCC 6633, respectively; j=5, 8, 9, where j=5- corresponds to mono+di nucleotide compositions and j=8,9 corresponds to SF: 8 – corresponds to genome similarity index dDDH and 9 – corresponds to index relied on similarity between DNA gyrase genes.

RF -Random Forest; SVM – LibSVM; KN – K-nearest neighbours; RA – RealAdaBoost; MP – MultilayerPerceptron; DM – Dl4jMlpClassifier

BAC and ACC was evaluated using 10-fold cross-validation.
